# Formula alters preterm infant gut microbiota and increases its antibiotic resistance load

**DOI:** 10.1101/782441

**Authors:** Katariina Pärnänen, Jenni Hultman, Reetta Satokari, Samuli Rautava, Regina Lamendella, Justin Wright, Christopher J. McLimans, Shannon L. Kelleher, Marko Virta

## Abstract

Infants are at a high risk of acquiring infections caused by antibiotic resistant bacterial strains. Antibiotic resistance gene (ARG) load is typically higher in newborns than in adults, but it is unknown which factors besides antibiotic treatment affect the load. Our study demonstrates that inclusion of any formula in the newborn diet causes shifts in microbial community composition that result in higher ARG loads in formula-fed infants compared to infants not fed formula. The effect of formula was especially strong in premature newborns and newborns treated with antibiotics. Interestingly, antibiotics alone without formula did not have a detectable impact on the ARG load of the newborn gut. We also observed that formula-fed infants had enriched numbers of pathogenic species and were depleted in typical infant gut species such as *Bifidobacterium bifidum*. The results suggest infant feeding choices should include assessment of risks associated with elevated ARG abundance.

## Background

Antibiotic resistant pathogenic bacteria cause approximately 214,000 newborn deaths annually^1^. Mortality related to antibiotic resistant infections is likely to rise and is estimated to surpass mortality caused by cancer despite efforts to the limit use of antibiotics^2^. Because the global spread of multi-resistant bacteria causes mortality and morbidity in newborns, understanding factors which affect the resistance load in this vulnerable group is crucial.

Previously it has been thought that selection due to antibiotic use plays the largest role in the increase in resistant bacteria in healthcare settings. However, recent reports on global antibiotic resistance have shown that antibiotic resistance does not correlate with antibiotic use, presumably because antibiotic resistant bacterial strains have already evolved and become globally omnipresent^3^. Consequently, transmission and suitable environmental conditions instead of contemporaneous selection caused by antibiotics are likely to explain the bulk of the observed clinical resistance. Therefore, limiting antibiotic use alone will not be adequate for controlling antibiotic resistance^3^. This trend can also be observed in the antibiotic resistance load of the infant gut microbiota as infants have more antibiotic resistance genes (ARGs) than adults even without having been exposed to antibiotics^4–7^. Thus, other selective factors besides antibiotics seem to drive the enrichment of resistant bacteria in the infant gut. The ARGs found in the infant gut can be transmitted from the mother^6^ or acquired from the hospital^8–11^. Consequently, it is critical to find ways to reduce transmission and modulate the gut environment to be less favorable for resistant bacteria.

There is limited knowledge on other factors besides antibiotics that can influence the resistome, defined as the collection of ARGs^12^, found in the infant gut. Feeding type has been observed to cause shifts in the abundances of specific ARGs in the infant gut^6,13^. However, the magnitude of the impact of diet on the total resistance load particularly in comparison to antibiotic use is currently unknown. Birth mode, gestational age, infant age as well as maternal factors such as antibiotic have been indicated to be able to modulate infant gut microbiota composition^14^. However, it seems that in newborns, feeding type might have the strongest effect on the gut microbiota^4,15,16^.

We hypothesized that the preterm neonatal gut resistome changes in response to feeding type, and that the change in the microbiota and resistome might cause changes in the resistance load. To investigate this hypothesis, metagenomic data from fecal samples collected at the age of 7 to 36 days from 46 preterm infants born between 27 and 36 weeks of gestation coupled with extensive metadata were fed to a machine learning algorithm to leverage explanatory variables which affect the resistance load (Supplementary table 1). A model built based on the 46-infant cohort was tested and improved using an additional set of 697 publicly available newborn gut microbiota metagenomes collected from five studies^17,4,5,13,18^. Our results show that formula feeding increases antibiotic resistance load in the preterm population. This finding has clinical implications for deciding feeding type for newborns when mother’s own milk is not available.

## Results

### Description of the cohort

The infants in this study were selected from a larger set of infants born at Penn State Hershey Medical Center (PSHMC) or transferred to the PSHMC neonatal intensive care unit (NICU) within 72 h of birth between 26–37 weeks of gestation (n = 46). Full description of the metadata for the cohort is available in Supplementary Table 1. Twenty-five infants were fed a commercial infant formula (Neosure), 20 were fed mother’s milk with human milk fortifier (Similac), and five received only breast milk (mother or donor). Thirty of the infants received antibiotics. In cases where the infant received antibiotics, fecal samples were collected approximately two weeks after the termination of antibiotic treatment.

### Formula increases ARG and MGE abundance

Random forest machine learning algorithm was used to identify explanatory variables linked to ARG load. Random forest analysis revealed that diet was the most important explanatory variable, followed by gestational age, age of infant, infant infection status, antibiotic use history of the infant, and delivery method. The variables from random forests were used to build generalized linear models (GLMs) with gamma distribution and log_e_-links and chi-squared tests were used for testing nested models (α: *p* < 0.05).

Infants who were fed any formula had significantly higher abundances of ARGs normalized to 16S rRNA counts than infants who were fed only breast milk or who were supplemented with fortifier (gamma distributed generalized linear model (GLM) with loge link: fold change 4.9 and fold change 3.8 respectively; Tukey’s post hoc test: adjusted *p* < 0.01 for both; Figure 1a; Supplementary Table 2). A more specific encoding for diet including all the nutritional sources received by the infant as well as information of which feed was the most prominent source of nutrition was used for testing of the gamma distributed GLMs and the model confirmed the effect of formula in all feed combinations (gamma GLM: *p* < 0.05, Supplementary Tables 1 and 2). MGEs were significantly more abundant in the formula-fed infants compared to infants who were not fed formula (gamma GLM: *p* < 0.05, Figure 1b; Supplementary Table 2), indicating that the ARGs might be mobile. There were no significant differences between infants who received fortifier compared to infants who were fed only breast milk (Tukey’s post hoc test adjusted *p* > 0.05; Figure 1a; Supplementary Table 2), nor were there differences related to the type of fortifier used, either human milk or bovine milk derived (GLM, and Tukey’s post hoc test: adjusted *p* > 0.05; Supplementary Table 2).

**Figure 1:**
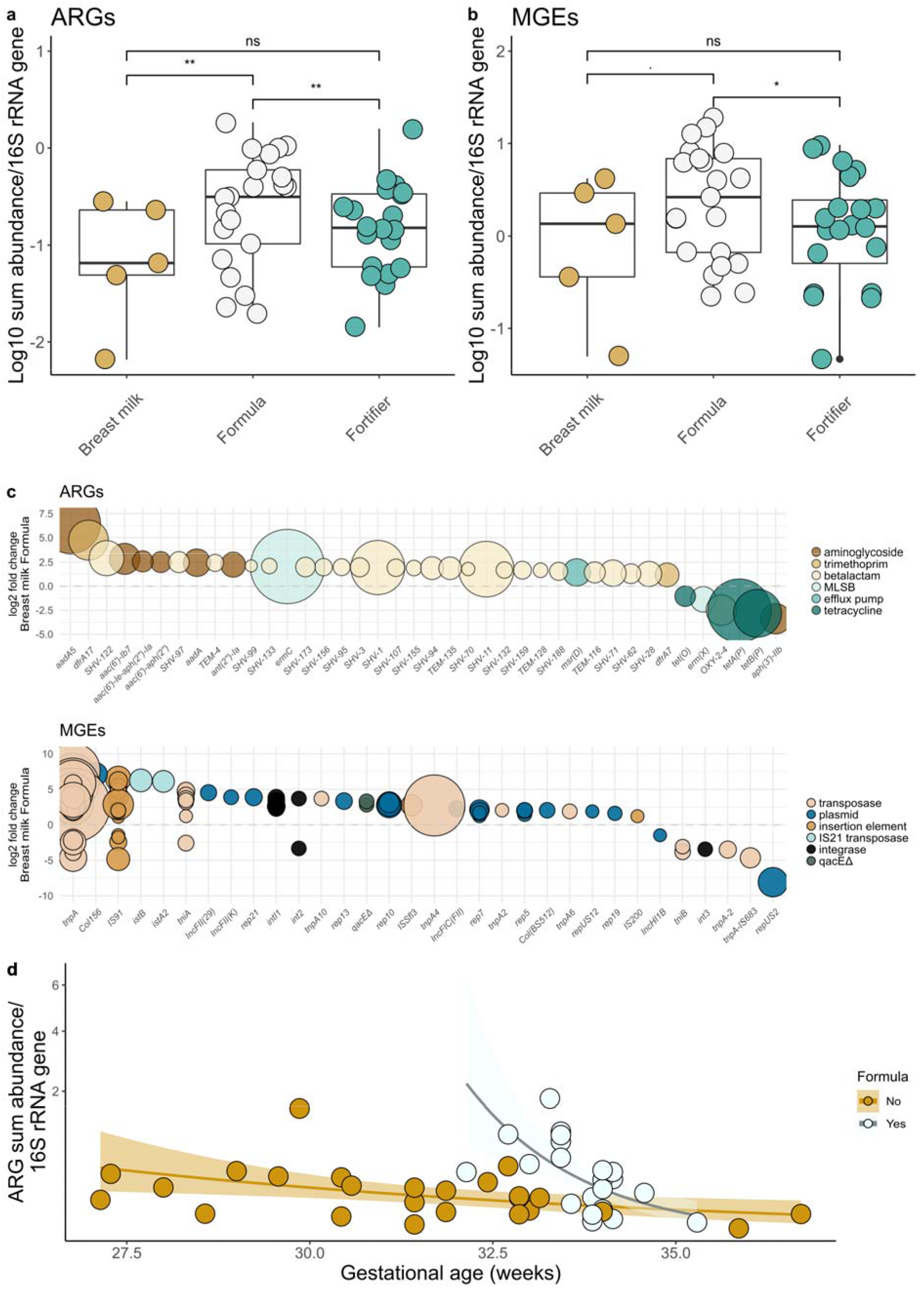
Total relative sum abundances of antibiotic resistance genes and mobile genetic elements by diet and gestational age. **a,** Relative ARG and **b,** Relative MGE sum abundances by diet. **c,** Differentially abundant ARGs and MGEs. **d,** Effect of gestational age and formula. In panels **a** and **b,** *y*-axis denotes the log_10_ transformed relative sum abundance. In **a** and **b** panels, significance levels are denoted as follows: ** = 0.001–0.01; * = 0.01–0.05;. = 0.05–0.1; ns = 0.1–1. In panel **c,** the size of the point represents the mean base level of the gene. In panel **d,** the x-axis represents gestational age in weeks. The x-axis has been square root transformed.

Many of the ARGs significantly enriched in infants fed formula were SHV-type beta lactamases (DESeQ2: adjusted *p* < 0.05; Figure 1c), which are typical beta lactamases found in *Klebsiella sp*. with some encoding for ESBL-phenotypes. The MGEs of all classes were enriched in formula-fed infants (DESeQ2: Figure 1c) including the integrase gene, *intl1*, which is part of the conserved region of class I integrons known to be able to give multidrug resistance phenotype to clinically relevant bacteria^19^.

The data included five twin pairs, and the generalized linear model (GLM) was also constructed without the twin pairs by including only the second-born infant, but this did not affect the GLM model estimates or significances. Twins were on average more similar in their microbial community composition, metabolic pathways as well as ARG and MGE composition, and more often had the same dominant species and shared strains with each other than with other infants (Supplementary Figure 1a–c and Strainphlan^20^ analysis).

### Gestational age, but not antibiotics, is linked to ARG load

Surprisingly, antibiotic treatment of neither the infant nor the mother was significantly linked to higher ARG load (gamma distributed GLM: fold change 1.2 and *p* = 0.68, and fold change 1.6 and *p* = 0.10, respectively). Gestational age and diet had a significant interaction (chi-squared test: *p* = 0.041). Gestational age was more strongly negatively correlated with the abundance of ARGs in the formula-fed infants (approximately 73 % lower abundance for one more week of gestation; gamma distributed GLM: p < 0.001; Figure 1d Supplementary Table 2) than in infants who were only fed breast milk with or without fortifier (approximately 23 % lower abundance for one more week of gestation; gamma distributed GLM: *p* < 0.001; Figure 1d; Supplementary Table 2). These result could indicate that formula feeding in preterm infants causes greater enrichment of ARG-carrying bacteria than in full-term infants.

### Formula increases intestinal *Enterobacteriaceae* in newborns

To investigate if we could observe differences in the abundances of microbial taxa know to commonly display antibiotic resistance, we compared abundances of *Enterobacteriaceae*, which are known to harbor several mobile ARGs in their genomes^21^, in infants with different diets. *Enterobacteriaceae* were more abundant in infants who were fed formula than in infants who were not fed formula (approximately three-fold higher abundance; quasibinomial GLMs, *p* < 0.05; Supplementary Table 2). *Enterobacteriaceae* was the dominant family in 29 of the 46 infants. These observations are in line with a previous report indicating that exposure to formula is associated with increased abundance of fecal enterobacteria^22^. In this study, similar to the ARG load, gestational age tended to be inversely correlated with *Enterobacteriaceae* abundance (fold change 22 % decrease / week of gestation; quasibinomial GLM:p < 0.1; Supplementary Table 2). Interestingly, *Enterobacteriaceae* have been hypothesized to be linked to the onset of necrotizing enterocolitis (NEC), and both lower gestational age and formula feeding are well-established risk factors for NEC^23–26^.

### Resistome correlates with community composition

The resistome and microbial community composition correlated significantly with each other (Mantel’s test for Horn-Morisita similarity index: *r* = 0.21, *p* = 0.001) in the 46-infant cohort. Moreover, there was a significant correlation between MGEs and the microbial species (Mantel’s test for Horn-Morisita similarity index: *r* = 0.27, *p* = 0.001).

The microbiota formed four clusters with *Veillonella, Klebsiella, Escherichia*, or Other (dominance by a genus which was dominant in only 1–2 infants) (Figure 2a). The metabolic genes were consistent between the microbial community composition and the metabolic pathways (PERMANOVA: false discovery rate adjusted *p* < 0.01, Figure 2b; Supplementary Table 3). ARGs clustered separately in *Escherichia*- and *Klebsiella*-dominated infants (PERMANOVA: false discovery rate adjusted *p* < 0.01; Figure 2c and 2d; Supplementary Table 3). MGEs clustered separately in infants dominated by Other and *Escherichia*, and *Klebsiella* and *Escherichia* (PERMANOVA: false discovery rate adjusted *p* < 0.05; Figure 2c and 2d; Supplementary Table 3).

**Figure 2:**
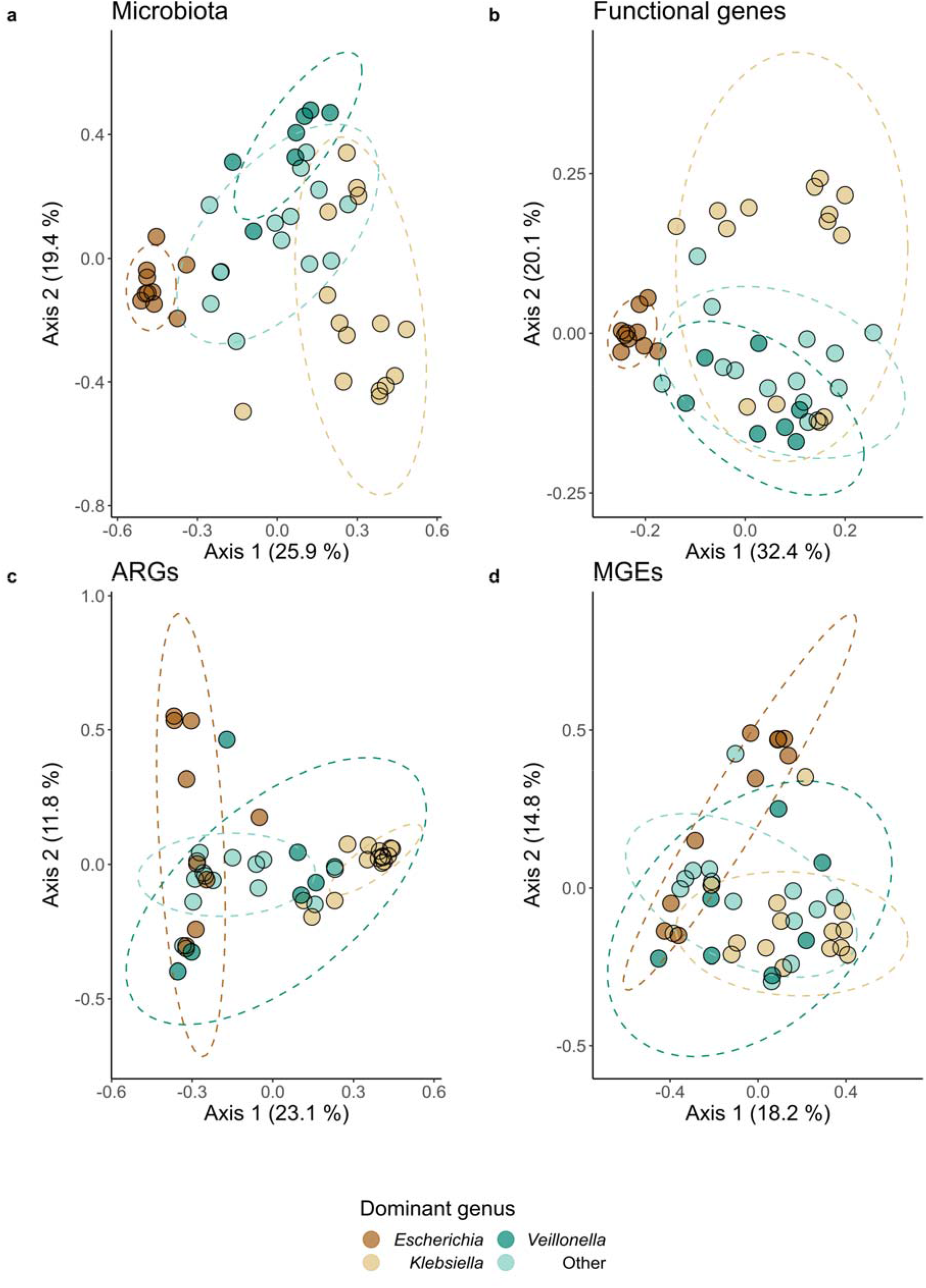
Clustering of microbial communities, functional gene pathways, ARGs and MGEs based on dominant species. **a,** Ordination of microbial communities using Metaphlan2. **b,** Ordination of metabolic genes using enzyme categories. **c,** Ordination of ARGs. **d,** Ordination of MGEs. Ordinations are performed using the Horn-Morisita similarity index with principal co-ordinate analysis (PCoA). Variation explained by axes 1 and 2 is denoted in in percentages. Confidence ellipses are draw for the three most common dominant genera using 90 % confidence level. *Klebsiella* was the most common dominant genus (n = 15) followed by *Escherichia* (n = 10) and *Veillonella* (n = 7).

The newborn gut communities were often dominated by one genus, which in some cases reached > 90 % relative abundance. Overall, the diversity of the microbial community in the preterm infant gut was very low with Shannon diversities ranging from 0.17 to 3.2 with a mean of 1.9 (Supplementary Figure 2a). There were no significant differences in the Shannon diversities between infants dominated by different genera (analysis of variance, AOV: false discovery rate adjusted *p* > 0.05; Supplementary Figure 2a and 2c–d). Metabolic gene diversity was higher in the *Veillonella-dominated* infants compared to other infants (AOV: false discovery rate adjusted *p* < 0.05; Supplementary Figure 2b).

### Formula is linked to higher ARG load in preterm infants

We sought to further investigate the impact of diet on the ARG load of infants by extending the analysis to include other metagenome sequencing projects in which feces of both full- and preterm newborn infants under the age of one month had been analyzed. We analyzed samples from four other metagenomic studies^17,4,5,13^ amounting to a total number of 242 newborn infants (age 0–36 days) including the original 46 infant cohort. In this meta-analysis cohort, 95 were fed only breast milk (own mother and/or donor milk) and 111 had formula in their diets. The remaining 36 were fed fortified breast milk and no formula. The metadata of the cohorts included in the meta-analysis are in the Supplementary Table 4.

After model selection, the best model based on chi squared tests for ARG load in infants of the meta-analysis cohorts included 16S rRNA gene counts, age of infant, gestational age and formula. The GLM results of the meta-analysis corroborated the observation made from the 46-infant cohort that formula feeding leads to higher relative abundances of ARGs compared to infants who are not fed formula (gamma distributed GLM: fold change 1.7, 95 % CI from 1.2 to 2.3, *p* < 0.01; Figure 3a; Supplementary Table 5). Gestational age and age alone had a significant effect on the relative ARG sum abundance, and the relative ARG abundances were 11 % / gestational week (95 % CI 7 % to 15 %) and 7 % / day of life (95 % CI 3 % to 10 %) (gamma distributed GLM: *p* < 0.001, Supplementary Table 5).

**Figure 3:**
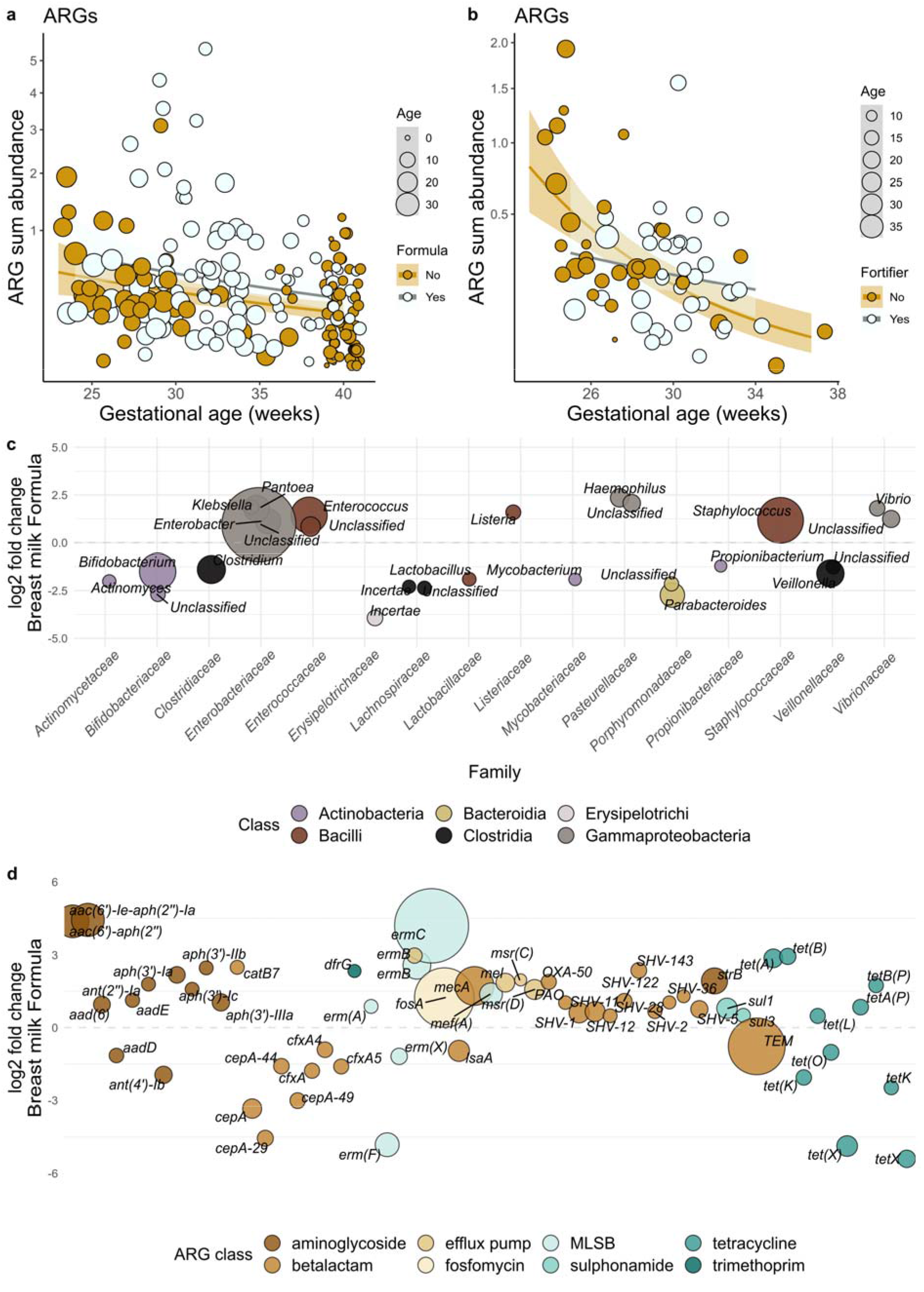
Effect of formula feeding on the resistome and microbiome of newborn infant gut in meta-analysis cohorts. **a,** Relative ARG sum abundance in relation to formula and gestational age. **b,** Relative ARG sum abundance fortifier use and gestational age. **c,** Significant differentially abundant genera, **d,** Significant differentially abundant ARGs between. In panels **a** and **b,** a regression line is fitted using a gamma distributed GLMs and the y-axes are square root transformed. In panel **c** and **d,** the colors describe the taxonomic and antibiotic classes. The y-axis depicts log_2_ fold changes observed between the groups. Size is proportional to the mean abundance, and only genera with a mean abundance > 10 and prevalence of > 10 % were included.

The addition of fortifier to breast milk did not significantly affect the sum abundance of ARGs (gamma distributed GLM: fold change 1.2, 95 % CI from 0.74 to 2.1, *p* = 0.42; Figure 3b, and fold-change 0.96, *p* = 0.91 for donor milk) in the meta-analysis. However, the analysis lacks statistical power necessary to draw definitive conclusions (power < 0.1).

### Formula has no detectable impact on full term ARG load

The interaction between gestational age and formula did not improve the model in the metaanalysis and the slopes with gestational age were similar for formula-fed and exclusively breast milk fed infants (chi-squared test: *p* > 0.05; Figure 3a; Supplementary Table 5), unlike in the 46-infant cohort. When analyzed separately in full-term infants (n = 83), formula had an increasing trend on the abundance of ARGs / 16S rRNA gene copies (fold change = 1.22, *p* = 0.43), but the effect was not significant despite being significant in preterm infants. Notwithstanding the fact that that we could not detect and effect of formula on the full-term ARG load, breast milk has been previously shown to decrease the abundance of specific ARGs in six-month old full-term infants^6^.

In order, to investigate further whether formula impacts ARG load in full term infants, we performed GLM analysis on a dataset of 501 full term infants aged less than one month from a study by Shao *et al.^18^* (See Supplementary Note 1 for list of ENA accession numbers). In this dataset 235 infants were fed only breast milk and 266 were fed formula with or without breastmilk. As in the full-term infants of the meta-analysis cohorts, the effect of formula on the ARG load was not significant. Among the clinical metadata available, delivery mode was the only significant explanatory variable that effected the ARG load in the Shao *et al*. cohort (24 % decrease in ARG sum / 16S rRNA genes in vaginally delivered infants compared to infant delivered by cesarean section).

### The effect of antibiotics on ARG load

Consistent with our dataset of 46 infants, antibiotic use, which was treated as a binary categorical variable (true or false), did not improve the model in the meta-analysis dataset (chi-squared test: *p* = 0.96, and gamma distributed GLM: fold change = 0.98, 95 % CI from 0.47 to 2.03, *p* = 0.96; Supplementary Table 5). However, gestational age and age were correlated with antibiotic use (Supplementary Table 5), which confounds our ability to fully differentiate between their effects on the ARG load. Nevertheless, age and gestational age were also significantly negatively correlated with ARG abundance in a subset of infants who all were treated with antibiotics (gamma distributed GLM: *p* < for both 0.01, Supplementary table 5). This confirms that the effect of gestational age and infant age observed in the model is not due to their correlation with antibiotic use.

The effect of formula on the ARG load did not reach statistical significance in the group of infants who were not treated with antibiotics (gamma distributed GLM: *p* > 0.05, Supplementary table 5), however the analysis lacked statistical power (power estimate < 0.5). A model with the interaction term between antibiotics and formula was fitted, and the interaction was significant (gamma distributed GLM: fold change 2.8 for infants with antibiotic treatment and formula compared to infants without formula and treatment, 95 % CI from 1.4 to 5.6, *p* = 0.0041), which might suggest that antibiotic treatment could increase susceptibility to colonization by ARG-carrying bacteria if an infant is formula-fed. However, it is not possible to infer whether this potential interaction is related to differences in gestational age in antibiotic treated versus non-treated infants (mean gestational age 29 weeks versus 38 weeks), since most preterm infants are routinely treated with antibiotics.

Antibiotic treatment did not affect species diversity in the meta-analysis dataset (fold change ≈ 1.0, *p* = 0.91 – 0.96, Supplementary table 6). Antibiotic treatment with specific antibiotics has been previously linked to decreased diversity in preterm infants^5,27^. Not differentiating between antibiotic classes can mask the effects that are caused by specific antibiotic classes.

### Formula feeding might increase pathogens possessing ARGs

In the meta-analysis cohort, there were significant differences in the microbial communities of infants receiving formula and infants on diets based exclusively on breast milk (PERMANOVA: *R^2^* = 0.0079, false discovery rate adjusted *p* = 0.041). This difference could not be captured in the 46- infant cohort. DESeQ2 analysis indicated that genera belonging to the obligate anaerobic families *Bifidobacteriaceae, Veillonellaceae, Clostridiaceae, Lachnospiraceae* and *Porphyromonadaceae* were depleted in the formula-fed infants of the meta-analysis cohort (DESeQ2: *p* < 0.05; Figure 3c) and in turn, the facultative aerobic families *Enterobacteriaceae, Staphyloccoccaceae* and *Enterococcaceae* were enriched.

The resistomes of formula-fed infants and infants fed breast milk clustered separately (PERMANOVA for Horn-Morisita similarity: *R^2^* = 0.011, false discovery rate adjusted *p* = 0.004) showing that besides increasing the prevalence of ARGs, formula feeding also affects the resistome composition. Many antibiotic resistance genes were enriched in the formula-fed infants (DESeQ2: *p* < 0.05; Figure 3d). The enriched genes included genes encoding beta-lactamases of the SHV type, which might confer extended spectrum beta lactamase (ESBL) phenotypes (DESeQ2: *p* < 0.05; Figure 3d) and the *mecA* gene encoding methicillin resistance in *Staphylococcus* species and the MLSB resistance gene, *ermC*, typically in found in *Staphylococcus aureus*. The SHV, *mecA* and *ermC genes* are all highly relevant ARGs in the neonatal intensive care units. Sulphonamide resistance genes of the *sul1* type were more abundant in formula-fed infants indicating the enrichment class 1 integrons which has been shown to often contribute to multidrug resistance for example in *Enterobacteriaceae*^19^.

Several pathogenic species including the facultative aerobes *Staphylococcus aureus, Staphyloccoccus epidermidis, Klebsiella pneumoniae, Klebsiella oxytoca* and the strict anaerobe *Clostridium difficile* were significantly enriched in the formula-fed infants compared to infants fed exclusively breast milk and typical infant associated bifidobacterial species, such as *Bifidobacterium longum*, as well as *Bacteroides* and *Parabacteroides*, were depleted (Metaphlan2^28^, DESeQ2: *p* < 0.001; Figure 4a, Supplementary Table 7). Interestingly, many of the depleted species were anaerobic while the enriched species tended to be facultative anaerobes. *Bifidobacterium* spp., in particular, are known to increase during breastfeeding as several species and strains are specialized in the utilization of human milk oligosaccharides^15^ and usually gain high abundances during breastfeeding in healthy full term infants shortly after birth^29^.

**Figure 4:**
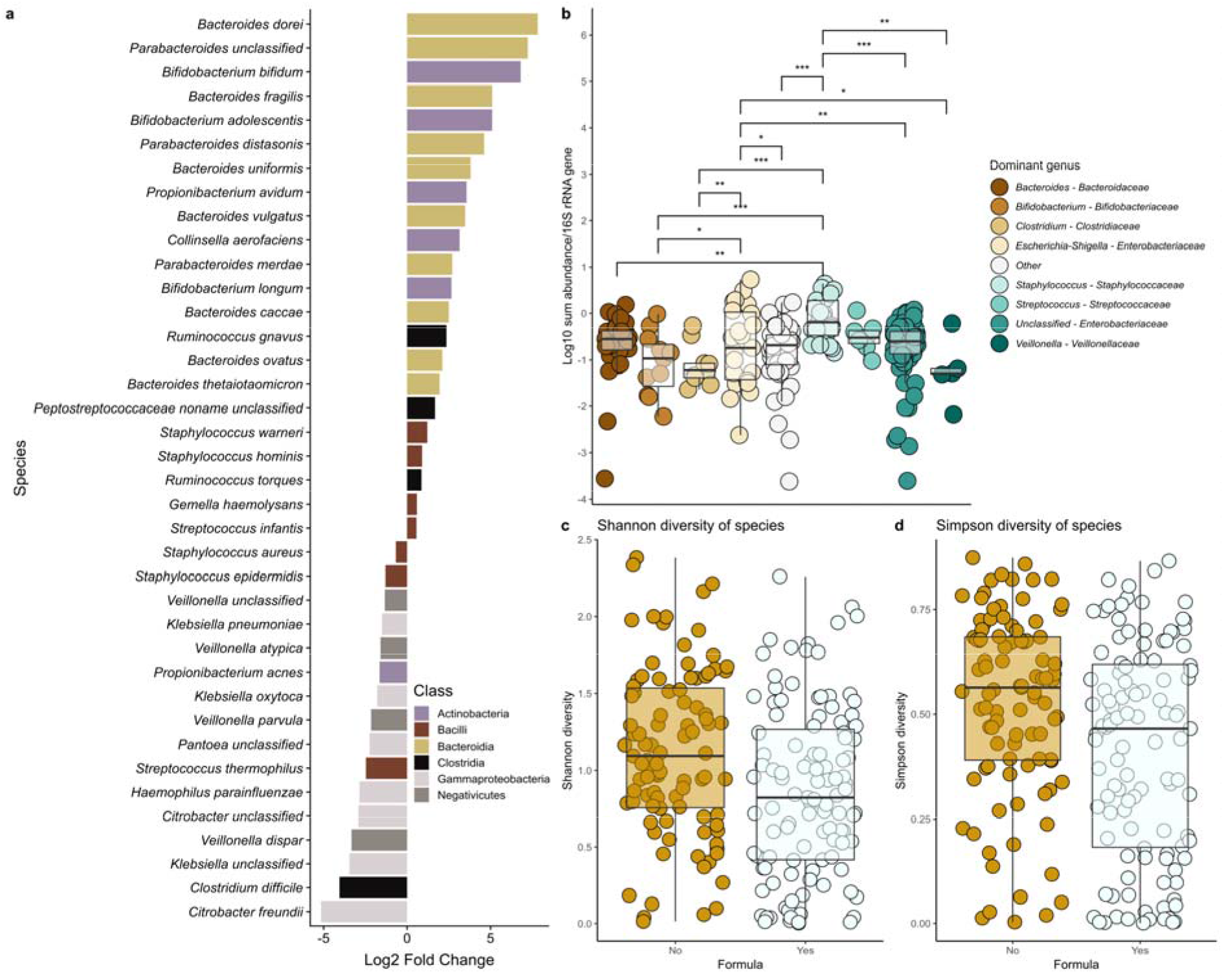
Microbial community changes linked to ARG abundances in meta-analysis cohorts. **a,** Species enriched or depleted in formula-fed infants. **b,** Relative ARG abundances in infants dominated by different genera. **c,** Shannon and **d,** Simpson (1-D) diversity index by formula. In panel **a,** negative/positive values reflect significant depletion/enrichment in formula-fed neonates. **In** panels **b, c** and **d,** the boxplot hinges represent 25 % and 75 % quantiles and center line the median. Notches are calculated with the formula median□±□1.58 × interquartile range / sqrt(n). Significance levels in panel **b** are denoted as follows: *** = 0–0.001; ** = 0.001–0.01; * = 0.01 – 0.05;. = 0.05–0.1.

We observed that the infants whose gut microbial communities were dominated by *Staphylococcus* had significantly more total ARGs / 16S rRNA copies than those dominated by *Bacteroides, Bifidobacterium, Clostridium, Streptococcus* and unclassified *Enterobacteriaceae*. Similarly, infants who had an *Escherichia/Shigella* dominance harbored more ARGs than those dominated by *Bifidobacterium* (gamma distributed GLM: *p* < 0.05; Figure 4b; Supplementary Table 8). The ARGs also clustered distinctly by the dominant genus, confirming that the microbial community composition is likely driving the differences in the resistome as well (PERMANOVA: *p* < 0.05; Supplementary Table 9).

Formula-fed infants exhibited significantly reduced diversity of microbial communities compared to infants who were not fed formula (Figures 4c and 4d; Supplementary Table 6), which has also been observed previously^22^. Since the formula-fed infants were also enriched in potentially pathogenic species and had more antibiotic resistance genes, it appears to be likely that potentially pathogenic ARG-carrying bacteria can reach dominance the formula-fed infant gut microbiota resulting in a simple community largely consisting of an ARG-carrying pathogen.

## Discussion

Our aim was to determine if newborn diet has an effect on the ARG load in the gut of the newborn preterm infant. We show that formula feeding increased ARG abundance compared to infants fed only breast milk particularly when formula-fed infants were also treated with antibiotics and/or premature. However, our study left open questions regarding the interaction between antibiotic treatment and gestational age in the effect of formula feeding on the ARG load. Our analysis suggests that either antibiotics are needed to aggravate the increase in ARG load in formula-fed infants, or that the more premature an infant is, the more strongly they are affected by formula. However, further studies are required to understand this relationship. Nevertheless, antibiotics alone did not have a detectable effect on the ARG load.

Formula-fed infants had increased abundances of *Enterobacteriaceae* as well as other potentially pathogenic bacteria. Also, ARGs that can, for example, confer methicillin resistance to *Staphyloccoccus aureus* and an ESBL phenotype to *Klebsiella pneumoniae* were increased compared to infants who were not fed formula. Interestingly, *Enterobacteriaceae* have been hypothesized to be linked to the onset of necrotizing enterocolitis (NEC), and prematurity and formula feeding are well-established risk factors for N EC^23–26^. The ARG load and resistome were linked to the microbial community structure, and taxa enriched in the formula-fed infants were correlated with higher resistance gene abundance. Our results indicate that in newborn, especially preterm infants, a diet containing exclusively breast milk or breast milk and breast milk fortifier reduces the antibiotic resistance gene load by modulating the microbial community to favor non-ARG carrying bacteria compared to a diet containing infant formula. This adds to the body of knowledge regarding the important health benefits of feeding breast milk particularly to preterm infants. Newborns and especially preterm infants are at risk of acquiring serious and life-threatening infections, and thus increased ARG loads in formula-fed preterm infants as well as the enrichment of potentially pathogenic bacteria represent matters of concern. Importantly, supplementing breast milk with fortifier was not associated with increased ARG abundance, which is reassuring in light of current guidelines concerning the nutritional management of preterm infants. The results suggest infant feeding choices should include the assessment of risks associated with elevated antibiotic resistance gene abundance linked to increased opportunistic pathogen prevalence in the preterm gut microbiota.

## Methods

### Ethical approval

The study was approved by the Institutional Review Board of Pennsylvania State University, USA (IRB #35925). Enrolment commenced on January 2014 and was concluded on July 2017. Inclusion criteria included the following: preterm infants born < 37 weeks of gestation who were admitted to the Penn State Hershey NICU or transferred to the PSHMC NICU within 72 h of birth. Exclusion criteria included the following: infants born > 37 week of gestation, or born with major congenital anomalies (heart, gastrointestinal, renal or respiratory tract), mothers known to use illicit drugs or abuse alcohol, or with a history of depression requiring long-term psychotropic medication. Written consent was obtained from all subjects. Metadata on feed history, clinical course, and NEC outcomes were collected electronically.

### Sample collection

Fecal material was collected into sterile microcentrifuge tubes approximately two weeks after prophylactic antibiotics had been discontinued and enteral feeds were initiated and frozen at −80 °C until analysis.

### DNA isolation and quantification

Fecal samples were collected and shipped to Wright Labs, LLC. Nucleic acid extractions were performed on approximately 0.25 g of each sample using the Qiagen DNeasy Powersoil DNA Isolation kit following the manufacturer’s instructions (Qiagen). The lysing step was performed using the Disruptor Genie cell disruptor (Scientific Industries). Genomic DNA was eluted in 50 μL of 10 mM Tris. Subsequent quantification was performed using the Qubit 2.0 Fluorometer (Life Technologies) using the double stranded DNA high sensitivity assay.

### DNA purification

The DNA was tested in standard 16S rRNA gene PCR to check for inhibitors before sequencing since some samples exhibited discoloration after DNA extraction. Samples which were discolored were observed to contain PCR inhibitors, and those samples were purified using the DNeasy PowerClean Cleanup Kit (Qiagen).

### Metagenomic sequencing

Metagenomic library preparation was performed using the Nextera XT library preparation kit (Illumina) using the standard protocol from the manufacturer. The library was paired-end sequenced using one run of NextSeq with an average of 10 million sequences per sample. Sequencing and library preparation were performed by the sequencing services of the Institute of Biology, University of Helsinki.

### Cohort description

The infants sequenced for this study were selected from a larger set of infants. Ethical permits for the study were approved by Penn State Hershey College of Medicine, USA. The inclusion criteria for the subjects were: infants born at Penn State Hershey Medical Center (PSHMC) or transferred to the PSHMC neonatal intensive care unit (NICU) within 72 h of birth between 26–37 weeks of gestation. Infants with 1) major congenital anomalies (heart, gastrointestinal, renal or respiratory tract) or 2) mothers known to use illicit drugs or alcohol, or with a history of depression requiring long-term psychotropic medications were excluded. Full description of the metadata for the cohort is available in Supplementary Table 1. The cohort included 46 premature infants. Fecal samples were collected between the ages of 7 and 36 days. Altogether, 29 of the infants were born by C-section and 17 were vaginally delivered. Twenty-five infants were fed a commercial infant formula (Neosure), 20 were fed mother’s milk with human milk fortifier (Similac), and five received only breast milk (mother or donor). Fortifier and formula were not given to the same infants, and therefore the subjects had either formula or fortifier as Supplementary feed. Thirty of the infants received antibiotics. In cases where the infant received antibiotics, fecal samples were collected approximately two weeks after the termination of antibiotic treatment.

### Metagenomic analysis

Quality control was performed using the FastQC^30^ and multiQC^31^ programs. The sequences were trimmed using Cutadapt^32^ version 1.10 with the options -m 1, -e 0.2, -O 15, and -q 20 to filter out adapters and low-quality reads. Filtered metagenomic sequence reads underwent subsequent microbial community profiling using the annotation software Metaphlan2^28^ version 2.6.0 with default settings. A merged abundance table was created using the Metaphlan2 ‘utils’ script. The merged abundance table was edited to only include taxa which were identified up to species level. 16S rRNA gene sequences were then identified, extracted and quantified using Metaxa2 version 2.6.0 in paired-end mode with default settings. The 16S rRNA gene sequence reads were classified using the mothur version v.1.40.5 ‘classify.seqs’ command with SILVAv. 123 as the reference database with the options cutoff = 60, probs = F and processors = 8. A custom Unix script was used to create an OTU table based on the classifications. *Escherichia coli* and *Klebsiella pneumoniae* strains were profiled using Strainphlan^33^ with the default settings. The ‘strainphlan.py’ command with the option -relaxed_parameters2 was used to create taxonomic trees of the strains in all samples. Metabolic genes were annotated using the Humann2^34^ pipeline with default settings and with enzyme categories (EC) for gene annotation and grouping.

Bowtie2^35^ was used for mapping reads to antibiotic resistance gene (ARG) and mobile genetic element (MGE) databases with the options -D 20, -R 3, -N 1, -L 20, and -i S, 1,0.50. Post annotation, SAMtools^36^ version 1.4 was utilized to filter and quantify observed ARG and MGE annotations within each sample. If both reads mapped to the same gene, the read was counted as one match, and if the reads mapped to different genes, both were counted as hits to the respective gene. We used the ResFinder^37^ database version 2.1 to search for acquired ARGs and a MGE database^6,38^ which includes genes related to or annotated as IS, ISCR, *intl1, int2, istA, istB, qacEdelta, tniA, tniB, tnpA*, and Tn*916* transposons. Plasmid related genes were searched for using the PlasmidFinder^39^ database.

### Meta-analysis cohort

To obtain more samples to study the variables affecting the ARG load in the newborn gut, a metaanalysis dataset was collected in addition to the original cohort of 46 infants. Literary searches were performed using the key words “metagenomics” AND “infant”, “preterm” OR “newborn” to find all available publications where newborn gut samples (with sampling on days 0 to 36 of life) had been shotgun metagenome sequenced using NextSeq of HiSeq (Illumina) with read lengths from 100 to 250 basepairs. Only sequencing datasets with 20 or more individual sampled infants were included. Other requirements were that data for gestational age, age at sampling, delivery mode and diet until the day of sampling as well as antibiotic treatment of the infant was available. After filtering out datasets which did not fulfill the qualifications, four datasets were included in the meta-analysis. The data was downloaded in fastq format, and in the cases where infants were sampled more than once, only the first of the samples in the time series was included. Only samples with more than 1,000 16S rRNA reads identified using Metaxa2^40^ were included in the meta-analysis. In total, the meta-analysis cohorts included 196 newborn samples. For datasets with twins, all models were run also without the firstborn twin to confirm that the estimates in the meta-analysis were not affected by including both twins.

The independence of the results related to the five different cohorts and sequencing platform and library preparation method combinations was confirmed by GLMs with gamma distribution as including these explanatory variables in the model did not improve the fit (results in Supplementary table 5). Also, the independence of the results related to the cohort used to choose explanatory variables to collect for the meta-analysis was verified by running the model without the cohort of 46 infants (results in Supplementary Table 5).

We postulated that formula given to the same infants as fortified breast milk would have similar effects as formula supplementation given to infants fed unfortified breast milk, but that fortifier fed infants might differ from infants receiving only breast milk. For the analysis of the effect of formula, we compared formula-fed infants to infants who were exclusively fed breast milk. To this end, we excluded all infants who were fed fortified breast milk but did not receive any formula, which allowed for the analysis in 206 newborn infants for comparisons between breast milk exclusive diets and formula containing diets. We separately analyzed the effect of breast milk supplemented with fortifier compared to an exclusively breast milk diet. In Bäckhed et al.^4^, gestational ages were between 39.3 and 41 weeks but were not reported separately for each infant, and an approximated gestational age of 40 weeks was used for all infants in this cohort. ARG annotation and quantification as well as Metaxa2^40^ and Metaphlan2^28^ community profiling were conducted for the obtained meta-analysis dataset as described above.

### Statistical analysis

All statistical analyses were performed in R version 3.6.1. Metaphlan2^28^, Metaxa2^40^ and Humann2^34^, annotation and ARG and MGE mapping results, taxonomy and metadata files were compiled into individual data objects in phyloseq^41^ version 1.28.0. All custom R codes are available in the Supplementary software file. The Bowtie2 counts for ARGs and MGEs were normalized to the length of each respective gene. Normalized gene counts were then further normalized to the number of bacterial 16S rRNA gene reads obtained from Metaxa2^40^ output divided by the length of the 16S rRNA gene. These normalization steps yield an approximation of the number of genes per 16S rRNA sequence for each resistance gene, while avoiding bias due to differential length of the resistance genes. The 16S rRNA gene was chosen for normalization instead of library size to account for variation in non-bacterial DNA content in the samples. The normalized values were used in all downstream analyses. All figures were plotted with ggplot2^42^ version 3.1.1.

Random forest analysis was performed using the caret^43^ package version 6.0-84 with training control using the ‘trainControl’ command with the option method = “cv”, and model fitting with ‘train’ command with the option method = “rf”, and the importance of the variable was computed using the ‘varImp’ command. The ARG sum relative abundance was used as response variable, and all variables in the metadata with diet being categorized as any formula, fortifier or breast milk, were used in the random forest algorithm (full list of variables available in Supplementary Table 1). Weight which correlated with both age of the infant as well as gestational age, and the type of maternal antibiotics, which had inadequate numbers of mothers receiving the same type of antibiotics to perform statistical analysis, were omitted from the analysis.

Different statistical model types were compared to select the best one for modeling the ARG / 16S rRNA gene ratio in the individuals of the 46-infant cohort. The data was checked for overdispersion, and as the variance was observed to be much larger than the mean, models with normality assumptions were not included as options. The data had positive continuous values and the responses seemed to be linear.Therefore, lognormal and gamma distributions were considered as possible distribution assumptions for the data. After comparing the two, gamma distributions were observed to capture more of the variance than log-transformed normal distributions, and the generalized linear model (GLM) with gamma distribution and a natural logarithmic link function was chosen. The GLM models were fitted with the ‘glm’ command using the option family = “gamma”(link = “log”). Model selection was performed using chi-squared tests by dropping out one variable or interaction term at a time and checking if the model performed significantly worse than the model with more explanatory factors. Diet was coded based on any formula (Formula), any fortifier (Fortifier) or no formula or fortifier (Breast milk).

The structure of the data in the cohorts included in the meta-analysis indicated that the relationships between the mean and variance were stable, with variance increasing with increasing mean, supporting the choice of a gamma distributed GLM similar to the cohort of 46 infants. Model selection was performed using all of the collected metadata variables (formula feeding, age, gestational age, delivery mode, and antibiotic treatment of the infant). The 16S rRNA gene counts in each sample calculated using Metaxa2^40^ outputs were also included in the model to account for potential effects of different library depths. The significance of the interaction between formula and gestational age was tested, since it had been significant in our cohort. The selection was performed using chi-squared tests, and the best model was selected by dropping out one variable at a time until the chi-squared test indicated that the model no longer improved.

Gamma distributed generalized linear mixed effect models with log links were explored to decide whether random effects caused by the study design as well as sequencing and library construction methods in the different meta-analysis cohorts should be included in the model. However, the models indicated that there was not enough variation between cohorts to warrant for random effects. Models with study as random effect resulted in singularity indicating that the random structure was too complicated. The random structure could not be simplified further. Thus, gamma distributed GLM without random effects with log link was chosen as the model type. For the analysis of the effect of formula, we compared formula-fed infants to infants who were exclusively breastfed. To this end, we excluded all infants who were fed fortified breastmilk but did not receive any formula, which allowed for the analysis of 206 newborn infants. We separately analyzed the effect of breastmilk supplemented with fortifier compared to an exclusively breastmilk diet. Power calculations to estimate effect sizes and sample sizes for a given effect size were done using the ImSupport package version 2.9.13 using the ‘modelPower’ and ‘modelEffectSizes’ commands.

Log linear models were also explored but they underperformed compared to the gamma distributed model and had AIC (Akaike information criterion, providing an estimate for the quality of statistical modes) values of around 700 compared to 40, and *R^2^* values of 0.16 compared to pseudo-*R^2^* values of the gamma GLMs of 0.6, with both metrics indicating worse performance.

Because Bäckhed at al. did not report antibiotic use for each infant individually, but only stated that 2 of the 100 infants were treated with antibiotics, these data were excluded from the model validation of the effect of antibiotics.

DESeq2^44^ version 1.24.0 was used to analyze differentially abundant taxa and genes. Analysis for Metaphlan2^28^ results was performed by multiplying the total 16S rRNA gene counts in each sample obtained from Metaxa2^40^ with the relative abundance values for each sample from Metaphlan2^28^. Metaxa2^40^ results were transformed by adding a pseudo count of one. Gene abundances were transformed by multiplying by 50,000 and rounded to integers resulting in normalized values, which consider variation in the lengths of the genes and the 16S rRNA gene counts in different samples. Using this transformation, the difference between a detected and an undetected gene is approximately 1.7-fold. A pseudo count of plus one was added to all samples to enable comparisons between detected and undetected genes. Several different options for transformation were tested but the results did not differ.

Principal coordinate analyses of the taxonomic profiles, metabolic pathways, ARGs and MGEs were performed using the ‘ordinate’ command from the phyloseq^41^ package. Distances between samples were calculated using the Horn-Morisita^45^ similarity index and oriented using principal co- oordinate analysis (PCoA). Permutational multivariate analysis of variance (PERMANOVA) between different groups was performed with adonis in the vegan^46^ package version 2.5-5 using 9999 permutations, and the resulting *p*-values were corrected with the Benjamini & Hochberg procedure for multiple testing using the ‘p.adjust’ command in R.

Distance matrices for species, ARGs and MGEs were calculated using the Horn-Morisita^45^ similarity index with the ‘vegdist’ command in the vegan package and compared to observe if there were correlations between microbial community and the ARGs and MGEs. Comparisons were performed using a Mantel’s test from the vegan^46^ package with the option method = ‘kendal’. Shannon and Simpson diversities for taxonomic profiles, ARGs and MGEs were calculated using the package vegan. Differences in the diversities were compared using analysis of variance combined with Tukey’s post hoc test.

The vegan^46^ package and Horn-Morisita similarity indices were used to calculate distance matrixes between twins and unrelated twins. Statistical analysis of the significances of observed differences between twin pairs and unrelated infants was performed by linear modeling with the ‘lm’ function in base R (see custom function ‘lmp’ in Supplementary software file). Permutations on how often a smaller *p*-value was achieved with randomized data as opposed to the assigned two groups compared was used to calculate a corrected p-value, with 9999 permutations used for the analysis.

## Acknowledgements

The research was supported by Academy of Finland. K.P. received funding from the MBDP doctoral program at the University of Helsinki, Finland. S.L.K received funding from The Gerber Foundation. R.S. is funded by the Sigrid Juselius Foundation, Finland. CSC, IT Center for Science, Finland, is acknowledged for providing the computational resources for the study. Abigail Podany is acknowledged for sample collection. Lasse Ruokolainen and Otso Ovaskainen are acknowledged for statistical consulting.

## Author Contributions

K.P. contributed to laboratory work and conception of the idea for the study, performed the statistical and bioinformatic analyses as well as drafted the first version of the manuscript. J.H., R.S., M.V., S.L.K., G.L., J.W. and S.R. contributed to the interpretation of the results. J.H., M.V., R.L., J.W., and S.L.K., contributed to conceiving the idea for the study. J.W. and J.M. contributed to laboratory work. S.L.K. provided samples and clinical data. All authors contributed to manuscript revisions and have read and approved the final version of the manuscript.

## Data availability

The sequence data is available from NCBI under the accession number PRJNA532310, which will be released upon publication in a peer reviewed journal.

## Code availability

All custom codes will be made available at https://github.com/KatariinaParnanen upon publication in a peer reviewed journal.

