## Supplementary note 1 for "Formula alters preterm infant gut microbiota and increases its antibiotic resistance load"

List of ENA accession numbers of the Shao et al., study for samples used in the analysis of formula impact on ARG load in full-term infants

ERS3420791

ERS3420792

ERS3420795

ERS3420804

ERS3420805

ERS3420807

ERS3420809

ERS3420810

ERS3420812

ERS3420817

ERS3420820

ERS3420822

ERS3420825

ERS3420827

ERS3420828

ERS3420830

ERS3420832

ERS3420835

ERS3420836

ERS3420837

ERS3420838

ERS3420842

ERS3420844

ERS3420845

ERS3420847

ERS3420850

ERS3420854

ERS3420857

ERS3420860

ERS3420861

ERS3420864

ERS3420866

ERS3420868

ERS3420871

ERS3420873

ERS3420877

ERS3420881

ERS3420887

ERS3420889

ERS3420891

ERS3420895

ERS3420898

ERS3420902

ERS3420905

ERS3420908

ERS3420910

ERS3420913

ERS3420918

ERS3420921

ERS3420924

ERS3420926

ERS3420929

ERS3420935

ERS3420940

ERS3420942

ERS3420946

ERS3420949

ERS3420953

ERS3420957

ERS3420960

ERS3420963

ERS3420965

ERS3420967

ERS3420971

ERS3420976

ERS3420979

ERS3420986

ERS3420990

ERS3420993

ERS3420996

ERS3421000

ERS3421003

ERS3421007

ERS3421009

ERS3421019

ERS3421026

ERS3421029

ERS3421034

ERS3421036

ERS3421039

ERS3421043

ERS3421047

ERS3421051

ERS3421054

ERS3421059

ERS3421064

ERS3421070

ERS3421072

ERS3421075

ERS3421077

ERS3421081

ERS3421088

ERS3421094

ERS3421096

ERS3421100

ERS3421105

ERS3421108

ERS3421111

ERS3421115

ERS3421118

ERS3421122

ERS3421126

ERS3421131

ERS3421135

ERS3421142

ERS3421146

ERS3421150

ERS3421151

ERS3421153

ERS3421156

ERS3421157

ERS3421159

ERS3421163

ERS3421166

ERS3421168

ERS3421170

ERS3421173

ERS3421176

ERS3421181

ERS3421183

ERS3421184

ERS3421188

ERS3421607

ERS3421610

ERS3421612

ERS3421614

ERS3421617

ERS3421619

ERS3421623

ERS3421625

ERS3421628

ERS3421630

ERS3421633

ERS3421635

ERS3421638

ERS3421641

ERS3421645

ERS3421648

ERS3421650

ERS3421653

ERS3421655

ERS3421657

ERS3421664

ERS3421665

ERS3421669

ERS3421671

ERS3421673

ERS3421676

ERS3421680

ERS3421682

ERS3421685

ERS3421687

ERS3421689

ERS3421693

ERS3421697

ERS3421699

ERS3421702

ERS3421707

ERS3421715

ERS3421720

ERS3421723

ERS3421725

ERS3421728

ERS3421729

ERS3421732

ERS3421733

ERS3421736

ERS3421739

ERS3421742

ERS3421744

ERS3421747

ERS3421752

ERS3421753

ERS3421758

ERS3421759

ERS3421761

ERS3421766

ERS3421768

ERS3421771

ERS3421773

ERS3421779

ERS3421781

ERS3421785

ERS3421787

ERS3421789

ERS3421792

ERS3421795

ERS3421797

ERS3421798

ERS3421802

ERS3421804

ERS3421807

ERS3421813

ERS3421818

ERS3421826

ERS3421829

ERS3421833

ERS3421835

ERS3421838

ERS3421841

ERS3421844

ERS3421849

ERS3421853

ERS3421857

ERS3421861

ERS3421865

ERS3421869

ERS3421873

ERS3421878

ERS3421881

ERS3421887

ERS3421892

ERS3421894

ERS3421897

ERS3421902

ERS3421905

ERS3421911

ERS3421912

ERS3421915

ERS3421917

ERS3421923

ERS3421927

ERS3421929

ERS3421932

ERS3421939

ERS3421943

ERS3421946

ERS3421949

ERS3421951

ERS3421954

ERS3421958

ERS3421961

ERS3421968

ERS3421971

ERS3421974

ERS3421977

ERS3421978

ERS3421982

ERS3421985

ERS3421994

ERS3421996

ERS3421998

ERS3422002

ERS3422007

ERS3422011

ERS3422016

ERS3422019

ERS3422022

ERS3422025

ERS3422032

ERS3422035

ERS3422039

ERS3422047

ERS3422048

ERS3422052

ERS3422055

ERS3422059

ERS3422062

ERS3422063

ERS3422067

ERS3422070

ERS3422075

ERS3422098

ERS3422102

ERS3422107

ERS3422110

ERS3422112

ERS3422116

ERS3422118

ERS3422119

ERS3422122

ERS3422125

ERS3422129

ERS3422132

ERS3422135

ERS3422139

ERS3422142

ERS3422145

ERS3422156

ERS3422158

ERS3422163

ERS3422168

ERS3422171

ERS3422175

ERS3422177

ERS3422179

ERS3422185

ERS3422187

ERS3422191

ERS3422201

ERS3422208

ERS3422212

ERS3422218

ERS3422221

ERS3422223

ERS3422226

ERS3422231

ERS3422234

ERS3422238

ERS3422241

ERS3422243

ERS3422247

ERS3422249

ERS3422252

ERS3422260

ERS3422263

ERS3422264

ERS3422267

ERS3422268

ERS3422270

ERS3422272

ERS3422274

ERS3422276

ERS3422280

ERS3422281

ERS3422283

ERS3422287

ERS3422294

ERS3422299

ERS3422301

ERS3422303

ERS3422306

ERS3422308

ERS3422312

ERS3422313

ERS3422315

ERS3422318

ERS3422320

ERS3422321

ERS3422323

ERS3422326

ERS3422327

ERS3422331

ERS3422332

ERS3422338

ERS3422340

ERS3422342

ERS3422344

ERS3422348

ERS3422350

ERS3422352

ERS3422354

ERS3422356

ERS3422358

ERS3422360

ERS3422363

ERS3422364

ERS3422366

ERS3422368

ERS3422372

ERS3422374

ERS3422376

ERS3422380

ERS3422382

ERS3422386

ERS3422387

ERS3422388

ERS3422390

ERS3422391

ERS3422392

ERS3422393

ERS3422394

ERS3422395

ERS3422396

ERS3422397

ERS3422399

ERS3422401

ERS3422403

ERS3422405

ERS3422407

ERS3422408

ERS3422409

ERS3422410

ERS3422411

ERS3422413

ERS3422414

ERS3422416

ERS3422421

ERS3422425

ERS3422431

ERS3422434

ERS3422437

ERS3422440

ERS3422444

ERS3422445

ERS3422447

ERS3422452

ERS3422456

ERS3422459

ERS3422461

ERS3422463

ERS3422468

ERS3422472

ERS3422473

ERS3422475

ERS3422482

ERS3422484

ERS3422488

ERS3422491

ERS3422497

ERS3422498

ERS3422505

ERS3422507

ERS3422509

ERS3422512

ERS3422514

ERS3422520

ERS3422522

ERS3422527

ERS3422529

ERS3422531

ERS3422534

ERS3422536

ERS3422538

ERS3422541

ERS3422543

ERS3422545

ERS3422548

ERS3422553

ERS3422556

ERS3422564

ERS3422571

ERS3422574

ERS3422577

ERS3422580

ERS3422583

ERS3422588

ERS3422591

ERS3422596

ERS3422602

ERS3422605

ERS3422608

ERS3422611

ERS3422614

ERS3422616

ERS3422622

ERS3422625

ERS3422630

ERS3422632

ERS3422637

ERS3422639

ERS3422642

ERS3422650

ERS3422655

ERS3422657

ERS3422664

ERS3422674

ERS3422678

ERS3422682

ERS3422684

ERS3422691

ERS3422696

ERS3422701

ERS3422704

ERS3422710

ERS3422711

ERS3422716

ERS3422720

ERS3422725

ERS3422728

ERS3422731

ERS3422735

ERS3422737

ERS3422738

ERS3422743

ERS3422748

ERS3422749

ERS3422751

ERS3422755

ERS3422759

ERS3422761

ERS3422766

ERS3422769

ERS3422772

ERS3422775

ERS3422777

ERS3422779

ERS3422783

ERS3422786

ERS3422788

ERS3422791

ERS3422792

ERS3422794

ERS3422796

ERS3422799

ERS3422802

ERS3422804

ERS3422807

ERS3422809

ERS3422811

ERS3422813

ERS3422815

ERS3422817

ERS3422818

ERS3422820

ERS3422823

ERS3422825

ERS3422827

ERS3422829

ERS3422830

ERS3422832
