## Supplementary figures for "Formula alters preterm infant gut microbiota and increases its antibiotic resistance load"

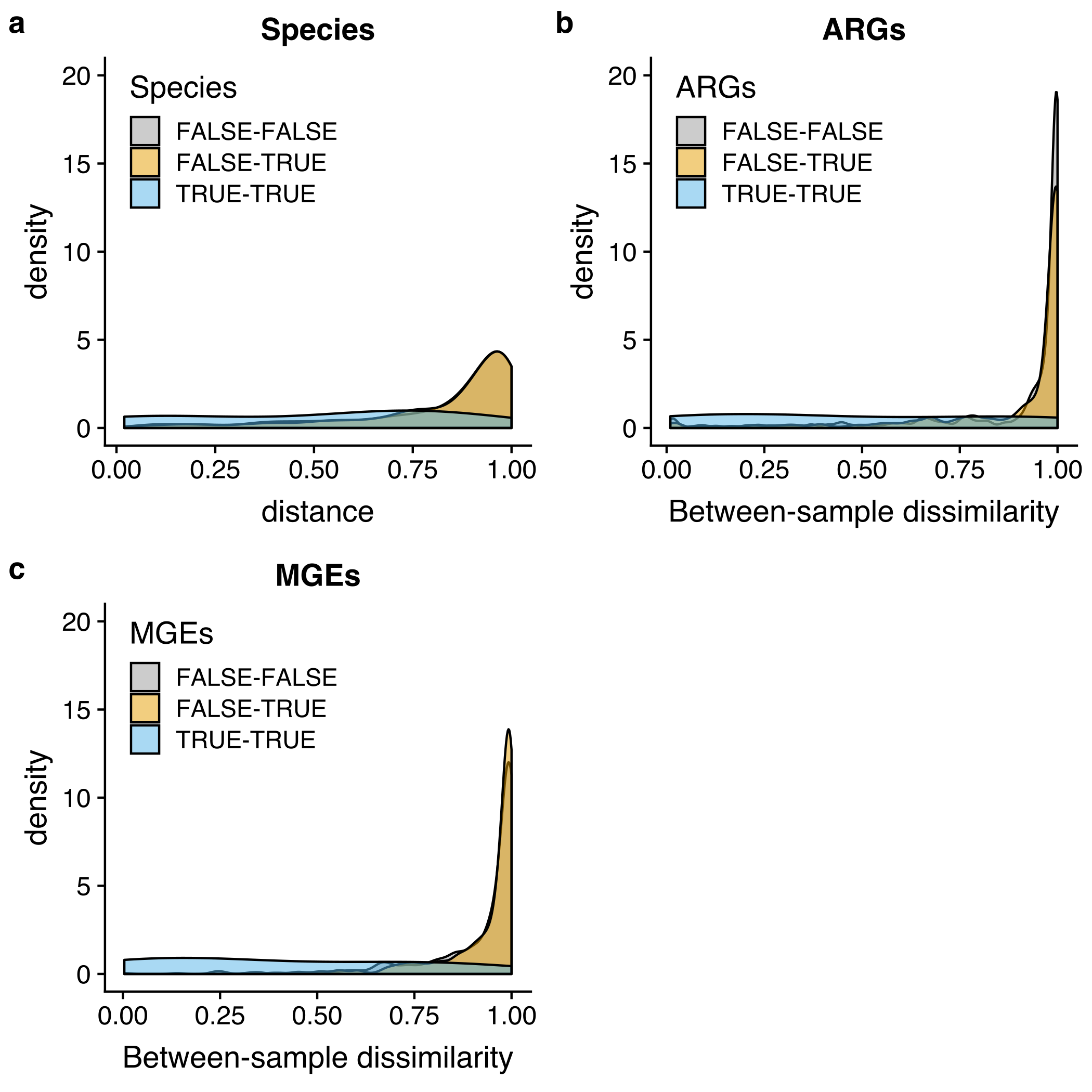

**Supplementary Figure 1: Resistome, mobilome and microbiota similarity between twins.**

**a**, Species similarity. **b**, ARG similarity. **c**, MGE similarity. The density plot depicts where comparisons between infant pairs are located on the dissimilarity scale. Pairings are denoted by color as follows: FALSE-FALSE indicates non-twin infant pairs with different diets; FALSE-TRUE indicates twin pairs with different diets; and TRUE-TRUE indicates twin pairs with same diets. Diet is classified into two categories based on the presence or absence of formula in the diet. The comparison for the similarity between twins is performed between non-twin pairs and twin pairs with same diets. Twins with same diet were consistently significantly more similar in their microbiota composition, ARGs and MGEs (ANOVA: *p* < 0.05). The higher the density is at a given dissimilarity value, the more pairwise comparisons have the given dissimilarity value. The density of the samples is plotted on the *y*-axis and the *x*-axis depicts the between-sample Horn-Morisita similarity index of species, ARGs and MGEs shared between sample twins

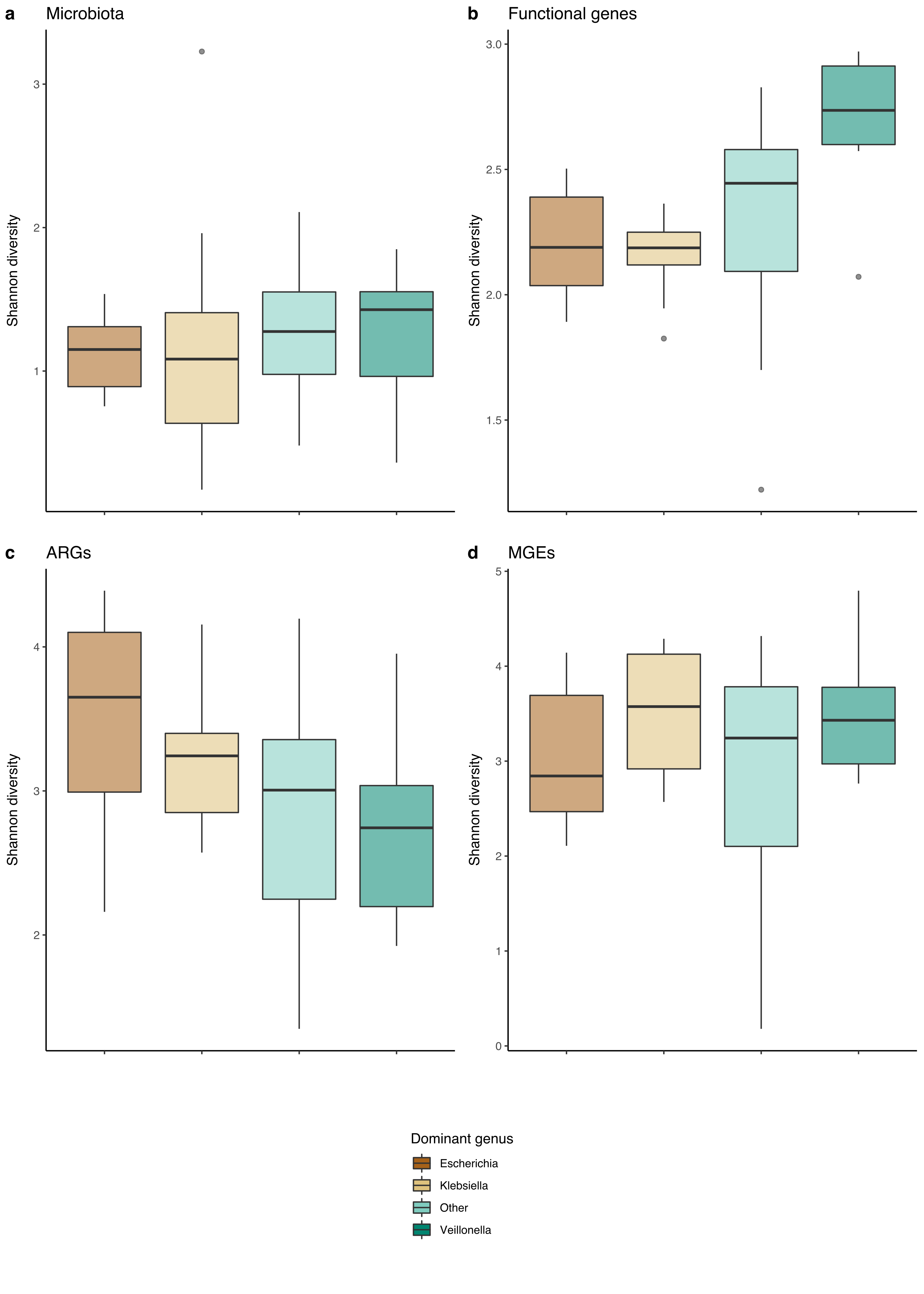

**Supplementary Figure 2: Diversity of microbial communities, functional gene pathways, ARGs and MGEs based on dominant species.**

**a**, Shannon diversities of species using Metaphlan2. **b**, Metabolic gene diversity. **c**, ARG diversity. **d**, MGE diversity. Diversities are computed using the Shannon diversity index. Samples are colored based on the most abundant genus in each sample. In boxplots **a** to **d**, the lower hinge represents the 25 % quantile, upper hinge the 75 % quantile and center line the median. Notches are calculated with the formula median ± 1.58 × interquartile range / sqrt(n). *Klebsiella* was the most commonly dominant genus (n = 15), followed by *Escherichia* (n = 10) and *Veillonella* (n = 7).
